# Fitness Functions for RNA Structure Design

**DOI:** 10.1101/2022.06.16.496369

**Authors:** Max Ward, Eliot Courtney, Elena Rivas

## Abstract

An RNA design algorithm takes a target RNA structure and finds a sequence that folds into that structure. This is fundamentally important for engineering therapeutics using RNA. Computational RNA design algorithms are guided by fitness functions, but not much research has been done on the merits of these functions. We survey current RNA design approaches with a particular focus on the fitness functions used. We experimentally compare the most widely used fitness functions in RNA design algorithms on both synthetic and natural sequences. It has been almost 20 years since the last comparison was published, and we find similar results with a major new result: maximizing probability outperforms minimizing ensemble defect. The probability is the likelihood of a structure at equilibrium and the ensemble defect is the weighted average number of incorrect positions in the ensemble. Also, we observe that many recently published approaches minimize structure distance to the minimum free energy prediction, which we find to be a poor fitness function.

## 1 Introduction

Ribonucleic Acid (RNA) is a versatile molecule [1]. It is fundamental to life [2] and has a breadth of roles including transcription and translation [3], catalyzing reactions [4], gene regulation [5], maintaining telomeres [6], and beyond. The functions of these RNAs are often governed by their structure [7]. Additionally, synthetic RNAs can easily be constructed [8]. A consequence of these properties is that RNA is a popular tool for bioengineering therapeutics (including mRNA vaccines [9]) and biomachines [10, 11, 12].

RNAs have the quality that fast and (relative to other molecules) accurate algorithms exist to predict their structures *in silico* [13, 14, 15, 16, 17]. Since structure usually determines function in biology, a multitude of attempts have been made to develop algorithms to automatically design RNAs with specific structures [18]. We refer to this as the *RNA design* problem. Algorithmic RNA design dates back to at least the early 90s [19]. These algorithms have led to some promising, practical successes [20, 21, 22, 23]. However, reliably solving RNA design puzzles is an open problem. No algorithm has solved the entire EteRNA100, a widely used RNA design benchmark that was hand-crafted by humans [24]. Recent results suggest the RNA design problem is NP-hard by proving that the most general version of the problem is hard [25] as well as a simplified model [26, 27].

Many techniques have been applied to the RNA design problem. We give a non-exhaustive sample for context. The first method was likely RNAinverse [19], which applied an adaptive random walk. Later methods used stochastic local search [28] with seeding heuristics [29], genetic algorithms [30, 31, 32], constraint programming [33, 34], simulated annealing [35], hierarchical decomposition [36, 37], Monte Carlo methods [38, 39, 40], and ant colony optimisation [41]. The EteRNA project has turned RNA design into a game in which solutions to RNA design puzzles can be crowd sourced from human players [42]. Recently, deep learning has also been applied to the problem in several ways including learning from human solutions [43], and by applying reinforcement learning [44, 45].

Each of these RNA design algorithms generally have three components,

1. A computational *model* (encompassing multiple components [17]), that can predict the structure of an RNA given its sequence.
2. A *fitness function*, which determines the quality of a potential solution with reference to a target structure.
3. A *search algorithm*, which explores the sequence space to find a desirable solution under the fitness function. The choice of possible search algorithms is often limited by the choice of fitness function.

The thermodynamic “nearest neighbor” model developed by Turner, Mathews, Tinoco and many others [46] is ubiquitous in the RNA design literature. The implementation of the nearest neighbor model in the ViennaRNA package was used in our work. This is the most widely used implementation [13]. However, it should be noted that other models exist and may offer new options for different fitness functions and search algorithms [17].

Generally speaking, design algorithms require a *fitness function* to guide their search. This is a measure of how promising an intermediate solution is. The fitness functions used are almost as numerous as the algorithms. However, most algorithms use fitness functions that are a variant of one of the four main types. These will be defined formally in Section 1.2. They comprise *structure distance minimization, free energy minimization, probability maximization*, and *ensemble defect minimization*.

Our aims are to summarize information about fitness functions for RNA design, and to provide an objective analysis of them. The latter is achieved by experiments. We use a fitness function agnostic search algorithm to do a fair benchmark. Both our aims are relevant to future works developing computational approaches to RNA design. The only similar work we are aware of is by Dirks *et al*. in 2004 [47]. We significantly expand and modernize the methods used, and find new results that differ somewhat from those of Dirks *et al*. We find that probability maximization performs better than any other fitness function, which differs from the previous result. This has implications for algorithms for RNA design, many of which currently use other fitness functions.

### 1.1 The Computational RNA Design Problem

Before we begin our discussion of fitness functions, we must understand the rules of the game: what is RNA design?

An RNA is represented by a string *p* ∈ {`A`, `U`, `G`, `C}`^∗^. The four characters in the alphabet correspond to the nucleotides that form RNAs, and the string represents the chain of nucleotides which constitute the primary structure of the RNA. We refer to this string *p* as the *sequence*. A secondary structure *s* is a (possibly empty) set of pairings between the indexes of *p* such that an index appears in *s* at most once and there are no two elements (*i, j*) ∈ *s* and (*k, l*) ∈ *s* such that *i* < *k* < *j* < *l*. In short, *s* is a properly nested set of pairings in *p*. In nature, some RNA structures contain improperly nested pairings called pseudoknots. Since the computational model we use does not allow these, we do not consider them. Henceforth we elide *secondary structure* to just *structure* for brevity.

Call *S*(*p*) the set of all possible structures for a sequence *p*. Define a *model m*(*p, s*) as a function that assigns every *s* ∈ *S*(*p*) a score. Since the structure represents bonds that form as the RNA sequence folds, we call the process of finding the optimal structure *folding* and define Fold(*m, p*) = arg min_*s*∈*S*(*p*)_ *m*(*p, s*). Ties for the arg min are broken arbitrarily. The solution to Fold(*m, p*) is the best guess of the true structure of the RNA under the model. Assuming the widely used thermodynamic model [46], Fold(*m, p*) can be computed in *O*(|*p*|^3^) time [16, 48].

The algorithmic RNA design problem is essentially to invert the folding problem. The folding problem takes a sequence *p* and finds an optimal structure *s*. The design problem inverts this relationship by taking a structure *s* and finding a sequence *p* that folds into *s*. Formally, we are given a structure *s* and a model *m*. Define the inverse folding function Fold^−1^(*m, s*) = {*p* ∈ {`A`, `U`, `G`, `C}`^|*s*|^ | Fold(*m, p*) = *s*}. Any sequence *p* ∈ Fold^−1^(*m, s*) is a solution to the RNA design problem.

Sometimes, there can be ties for the optimal fold Fold(*m, p*). This happens when there are multiple structures with the minimum score. This issue is often overlooked in the literature. There are many published results where a folding algorithm is run and whatever arbitrary structure it produces in the case of ties is taken—a non-exhaustive list: [41, 38, 42, 18]. There are two ways to address the issue. We say that a sequence *p* is a correct design for a structure *s* when it is tied with the optimal fold: *m*(*p, s*) = *m*(*p*, Fold(*m, p*)). Alternatively, we can say that *p* is a correct design iff Fold(*m, p*) = *s* and there are no ties. We adopt the latter tie-breaking strategy because it is more stringent and should lead to more accurate designs *in vivo*. If there are ties for an optimal structure, it means the model cannot determine which is the most likely. An ideal design should ensure the target structure is non-ambiguously selected under the model.

### 1.2 Fitness Functions

Success on the algorithmic RNA design problem as specified in Section 1.1 is binary. Either a design is correct, or it is not. This is not directly usable for most search algorithms, which need to be able to make incremental progress toward a solution. This is why fitness functions are important. Their use is to estimate the effectiveness of intermediate sequence solutions towards a target structure. Next, we describe the typical fitness functions used in RNA design.

#### 1.2.1 Structure Distance Minimization

The key idea in structure distance minimization is to make Fold(*p, m*) as similar to the target structure as possible. Given a distance function *d*(*s*_1_, *s*_2_) between two structures, the structure distance for a given potential solution sequence *p* and target structure *t* is *d*(*t*, Fold(*p, m*)). Minimizing the structure distance can lead to a correct design. This category of fitness function seems to be the most widely used [30, 19, 38, 35, 39, 45, 28].

It is important to choose a good distance function. In our experiments, we use two that are widely used in the literature. The first is base pair distance as implemented in the ViennaRNA software package [13]. Base pair distance is the number of base pairs that occur in exactly one of the two compared structures. The second is the number of indexes in *p* where the structure differs from the target, defined as the number of incorrect nucleotides in [47].

A weakness of using structure distance is that it is unclear what to do in the case of ties for the optimal fold. The typical approach in the literature seems to be not to address this edge case and accept the arbitrary result of Fold for structure distance comparison. We try several methods of dealing with ties including using the average distance and minimum distance. Another weakness is that structure distance myopically considers only the results of Fold while ignoring all other structures. This can make structure distance a poor guide when the target structure is close to optimal, but not an optimal fold. Due to these weaknesses, we expected that structure distance is not an effective fitness function. The expectation that structure distance is a poor metric may be considered controversial, since it is widely used. However, the previous work on fitness functions by Dirks *et al*. found structure distance (they call it MFE satisfaction) to have middling efficacy [47].

#### 1.2.2 Free Energy Minimization

Most existing RNA design algorithms use a thermodynamic nearest neighbour model. This model assigns each structure *s* ∈ *S*(*p*) a free energy score. Lower free energy structures are more likely at equilibrium and therefore more likely to be the true structure. Free energies can be used directly for RNA design. The utility of a potential design *p* for a target structure *t* can be defined as *m*(*p, t*). It seems natural that we want to select designs that minimize the free energy of our target structure. This fitness function is often used in conjunction with other metrics [30, 29, 38], but it has been used independently too [47, 49].

Despite its intuitive appeal, free energy minimization has a major weakness when used for algorithmic RNA design. Consider two sequences *p*_1_ and *p*_2_ and a structure *s* ∈ *S*(*p*_1_) ∩ *S*(*p*_2_) where *m*(*p*_1_, *s*) > *m*(*p*_2_, *s*). It is possible that Fold(*m, p*_1_) = *s* and Fold(*m, p*_2_) ≠ *s*. In words, *s* may be more stable in *p*_2_ compared to *p*_1_, but not the most stable among all structures with respect to *p*_2_ while being the most stable over all structures for *p*_1_. This is because a “good” score in *S*(*p*_1_) may not be a “good” score in *S*(*p*_2_). In essence, we cannot sensibly compare scores drawn from different thermodynamic ensembles. Additionally, if free energy minimization was a perfectly effective fitness function, then either algorithmic RNA design would not be NP-hard or NP = P since a sequence with the minimum possible free energy for a structure can be found in polynomial time [29]. Due to this weakness, we expect that free energy minimization is not an effective fitness function. Dirks *et al*. previously found free energy minimization to be a relatively poor fitness function [47].

#### 1.2.3 Probability Maximization

The thermodynamic equilibrium partition function for RNAs under the thermodynamic model can be computed efficiently, requiring *O*(|*p*|^3^) time, the same as computing Fold(*m, p*) [50]. This allows us to compute the conditional probability **P**(*s* | *p*) of a structure *s* given a sequence *p*. This probability can be maximized and used as a fitness function for RNA design. This fitness function has been used for RNA design, [31, 47, 19] albeit less widely than some other fitness functions.

Using probability maximization is similar to free energy minimization, but it addresses many of the major shortcomings. Because the probabilities are normalized to the same scale, it is more sensible to compare two probabilities taken from different sequences. In this sense, probability can be thought of as a “better” free energy minimization. There is still a weakness, however. Suppose we are comparing two sequences *p*_1_ and *p*_2_ as potential designs for a target structure *t*. It is possible that *t* is the best ranked (by probability) for *p*_1_, but has a higher probability and a worse rank in *p*_2_. This can happen when the probability distributions looks dissimilar between sequences, for example *p*_1_ could have a flat distribution and *p*_2_ could have an exponential distribution.

This weakness is similar to the weakness of free energy minimization. However, it should have a lower negative impact, since it requires the distributions of structure probabilities for sequences to look dissimilar. For this reason, we expected that probability will be a better fitness function than free energy. Dirks *et al*. found that probability maximization was an effective fitness function [47].

#### 1.2.4 Ensemble Defect Minimization

Ensemble defect was originally introduced by Dirks *et al*. who found it to be the best fitness function they tested [47]. They called it “average number of incorrect nucleotides,” which describes what it measures. It was later called ensemble defect [36]. In essence, the ensemble defect of a sequence *p* with respect to a target structure *t* is the sum of a certain distance function over the ensemble of structures *S*(*p*) weighted by probability.

A structure *s* is a set of paired nucleotide locations. For convenience, let *s*_*i*_ = *j* if *i* and *j* are paired, and *s*_*i*_ = *i* if *i* is not paired to any nucleotide. Let our distance function be the average number of incorrect nucleotides between a structure *s* and a target structure *t* such that |*s*| = |*t*|, similar to a Hamming distance:

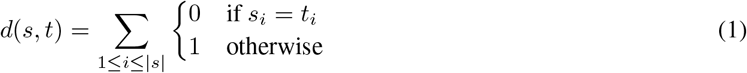

Define **P**(*s* | *p*) as the probability of the structure *s* given the sequence *p*. The ensemble defect of a sequence *p* for a target structure *t* is defined as:

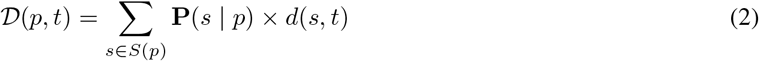

It is possible to compute 𝒟(*p, t*) in *O*(|*p*|^3^) time, making it no less efficient to compute than comparable fitness functions [47]. While not as widely used as structure distance minimization, ensemble defect is well known [36, 37, 33, 47, 51]. It appears to combine the strengths of probability maximization and structure distance minimization while avoiding the weaknesses of both. We expect that it will be the most effective fitness function.

## 2 Methods

We ran two different experiments. The first used synthetic RNAs, and the second used real RNAs. In the first, we used a fitness function agnostic search algorithm to test the performance of each fitness function type on synthetic RNAs. In the second, we took real RNA sequences with conserved structures and determined if the fitness functions predict that the true sequence is a good design for the true structure.

We used the ViennaRNA package 2.5.0 [13] as the model and the RNAfold program as the folding algorithm for all our experiments.

### 2.1 Synthetic RNAs

A search algorithm is needed to test the fitness functions. It must be compatible with all fitness functions. We used an Adaptive Random Walk (ARW), since it supports all fitness functions, is widely known, and is relatively simple. Dirks *et al*. [47] used an ARW, as does the RNAinverse program in ViennaRNA [19, 13]. The algorithm is described in Algorithm 1, and makes a sequence of mutations that maintaing base pairs and accepting only those that improve the fitness function score.

The ARW was run using each fitness function. ARW was run for 1000 steps on each target structure. A result sequence *p* was considered correct for a target structure *t* if it was the unique solution; that is it was the unique minimum free energy structure as judged by ViennaRNA [13]. If there were ties or if *p* did not fold into the target structure, then a result was considered incorrect. For each fitness function, we counted the number of correct and incorrect results. Better fitness functions should guide the ARW to good solutions more often, so the rate of correct results was used to measure fitness function efficacy.

#### Algorithm 1

Adaptive Random Walk

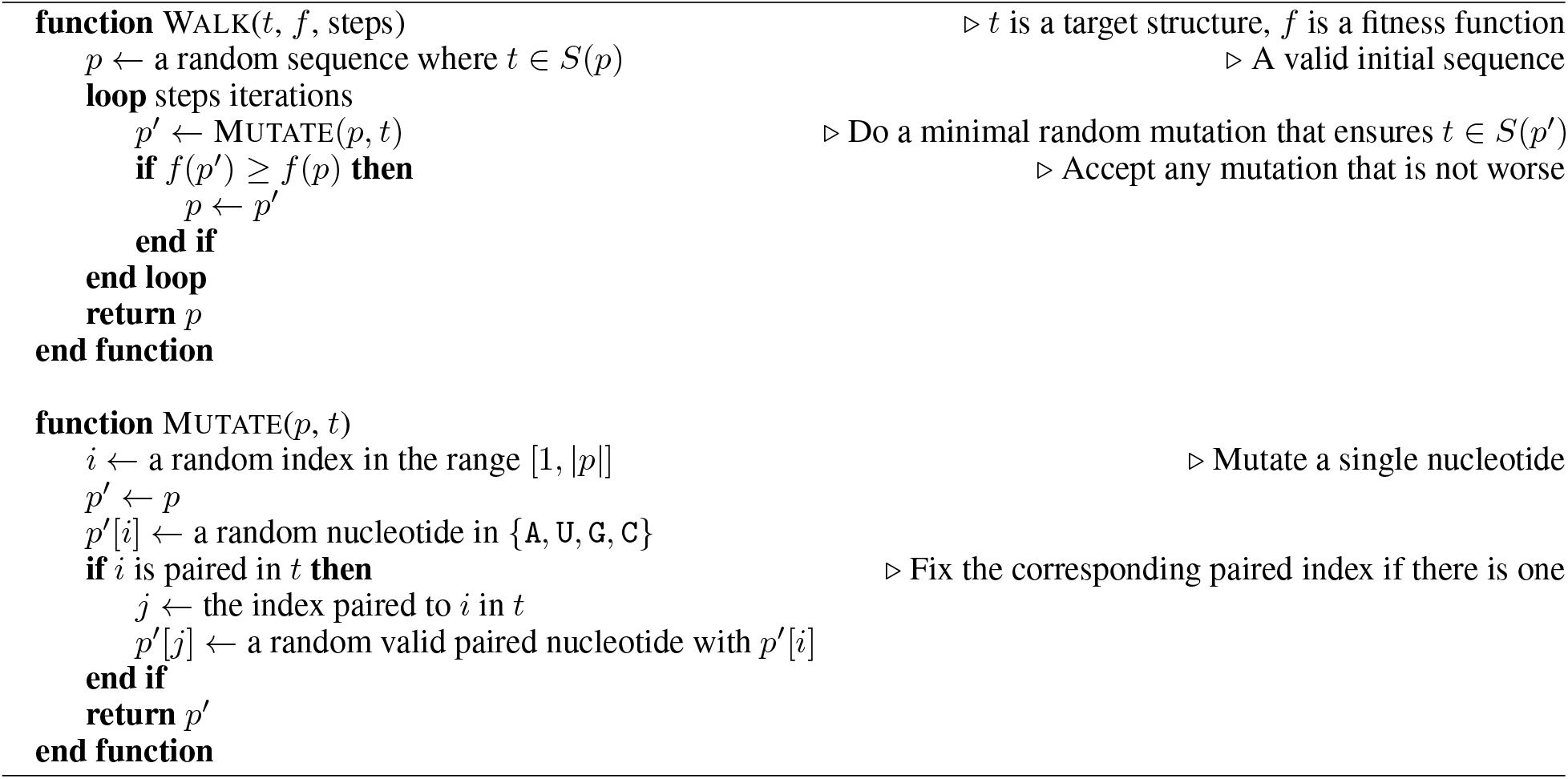

Two synthetic data sets were used. A “harder” data set and an “easier” data set. In both, target structures were generated the same way. First, sequences were generated by picking each nucleotide with equal probability. A target structure was picked randomly for each sequence from the set of structures within *T kcal/mol* of the minimum free energy. A different *T* value was picked in different data sets to adjust difficulty. We use this method because always picking a minimum free energy structure tended to generate structures that were easily solved by the ARW. The resulting set of target structures for each sequence was used as our testing data set. Wuchty’s algorithm [52] in ViennaRNA [13] was used to generate all suboptimal structures within the free energy window.

For the harder data set, 1600 structures of length 80 were generated with *T* = 5.0 *kcal/mol*. For the easier data set, 1600 structures of length 40 with *T* = 1.0 *kcal/mol* were generated. These values (1600, 80, 5, 40, 1) were picked before examining the performance results. Running the ARW for 1000 steps was an arbitrary parameter that was also chosen before examining performance results. Length 80 was picked because it was the largest length that ran in time on an AMD 5950X processor. Similarly, 40 was picked because it was half of 80. A window of *T* = 5.0 *kcal/mol* was chosen because 6.0 was too slow on our processor, and 1.0 was picked because it was significantly smaller than 5.0.

The previous work by Dirks *et al*. tested on 11 structures [47]. Of these, 9 were variants of the a tRNA inspired 3-way multiloop structure with varying stem lengths and numbers of unpaired nucleotides. Of the two remaining, one is a larger multiloop, and the other is a small pseudoknot. In contrast, we use 3200 sequences with a much greater degree of diversity due to our method of generation. In particular, we expect our harder set to contain much more difficult to solve structures. A major issues with the Dirks test set in addition to its small size is that all structures were solved by both probability and ensemble defect. We believe harder puzzles are needed to distinguish between these two.

We do not believe that our structure generation method has a bias toward producing structures that are easier for any particular fitness function, because the results using synthetic data are consistent across both data sets and with those from real RNAs. However, we could not find a way to ensure no bias in data generation for this or any other method except picking structures uniformly at random. Uniform random structure generation could not be used because we found it almost always generates structures no fitness function could solve.

#### 2.1.1 Structure Distance

We tried several variants of structure distance. We used “base pair distance” (BPD), which is a standard distance measure included in ViennaRNA [13] and defined in Section 1.2.1 as the number of base pairs that occur in exactly one of the two compared structures. We also used a structural Hamming distance (HD) defined the same as in Equation (1).

For both of these distance functions, we tried three different ways of dealing with ties for the minimum free energy structure. This issue is described in Section 1.2.1. We tried using the single arbitrary structure ViennaRNA returns; taking the average distance over all ties; and taking the minimum distance over all ties.

Both BPD and HD give distance values in the range from 0 to *n* for *n* nucleotides. Because our ARW implementation maximizes fitness values, we used *n* − *d*(*s*_1_, *s*_2_) as our fitness value where *d*(*s*_1_, *s*_2_) represents some distance function between the structures *s*_1_ and *s*_2_.

#### 2.1.2 Free Energy

The free energy as evaluated by the ViennaRNA package [13] was used. Because a lower free energy is more stable, and our ARW maximizes fitness, we use −Δ*G* as the fitness value, where Δ*G* is the free energy change.

#### 2.1.3 Probability

The probability P(*s* | *p*) for a structure *s* given a sequence *p* as evaluated by the ViennaRNA package [13] was used. This was achieved by computing the partition function for the sequence *p*. Probability was maximized directly by the ARW.

#### 2.1.4 Ensemble Defect

The normalized ensemble defect was calculated using the method provided in the ViennaRNA package [13]. This is the same as in Equation (2), but normalized by sequence length: 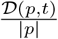. Because our ARW maximizes fitness, we optimized 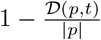.

### 2.2 Real RNAs

Many RNAs in nature have conserved structures. Consider a natural RNA sequence *p* with a known structure *s*. A fitness function *f* can be used to rank every structure in *S*(*p*) as a design for *p*. We can assign each structure *s*^*′*^ ∈ *S*(*p*) a score *f* (*p, s*^*′*^) then order *S*(*p*) according to these scores. A good fitness function is expected to rank the true structure *s* highly. Natural sequences have been “designed” through natural selection to fold into their structure. While the computational folding algorithm only approximates this, we do expect that the real structure for a natural sequence is seen as a good design. A subtlety worth emphasizing is that we are not comparing multiple sequences for a single target structure, as in Section 2.1. Instead, we are looking at all the structures for a single, natural sequence that has a single true structure.

Using natural sequences is important to fairly compare ensemble defect. It is claimed that ensemble defect is expected to make good predictions on natural sequences due minor base pairing fluctuations *in vivo* and the observed low ensemble defect of natural helices [47, 36, 37]. Despite this, the previous work on fitness functions by Dirks *et al*. [47] did not test on natural sequences, so we consider this analysis novel.

We used *ArchiveII*, a curated collection of 3948 natural RNA sequences with conserved structures [53]. For each, suboptimal folds were generated using the implementation of Wuchty’s algorithm [52] in ViennaRNA [13]. The suboptimal free energy window started at 0 *kcal/mol* and was increased in increments of 0.2 *kcal/mol* until at least 200 000 suboptimal structures were generated or the window exceeded 10 *kcal/mol*. The structures were ranked by their corresponding fitness function values. The rank of the RNA’s true structure was found in the ranked list of structures. If it was not found, the closest structure by base pair distance [13] was used. If the closest structure differed by more than 5% of the sequence length, the sequence was excluded from the experiment. A total of 1719 sequences were excluded this way. The 5% similarity threshold was picked because often the true structure does not appear in the suboptimal list (i.e. it contains a base pair not allowed in the nearest neighbour model), but a nearly identical structure does appear.

Probability, ensemble defect, and structure distance were considered in our experiment. We opted not to test free energy. In the special case of this experiment, free energy necessarily always has the same ranking as probability because the sequence is fixed. For structure distance, we considered only Hamming distance breaking ties for the minimum free energy by averaging as described in Section 2.1.1 because it was the best performing structure distance variant for synthetic RNA.

## 3 Results

### 3.1 Synthetic RNAs

Table 1 and Table 2 summarize the results from Section 2.1. The first table shows probability and ensemble defect with the strongest performances on the easier synthetic data set. Ensemble defect and probability perform similarly. In order, the performance seems to be free energy, structure distance, ensemble defect, probability. This is somewhat similar to the results found by Dirks *et al*. [47] who found that both probability and ensemble defect solved their entire test set.

**Table 1:** Results on synthetic structures of length 40. The “# Correct” is the number of correct solutions out of 1600. The “Correct Rate” is the ratio of the number of correct solutions and 1600. The “Unique Solver” column contains the number of structures for which a fitness function was the only fitness function to find a correct solution.

| Fitness Function | # Correct | Correct Rate | GC-percent | Unique Solver |
| --- | --- | --- | --- | --- |
| Structure Distance (BPD; arbitrary tie breaking) | 1269 | 0.79 | 0.50 | 1 |
| Structure Distance (BPD; average tie breaking) | 1419 | 0.89 | 0.50 | 1 |
| Structure Distance (BPD; minimum tie breaking) | 1135 | 0.71 | 0.50 | 0 |
| Structure Distance (HD; arbitrary tie breaking) | 1286 | 0.80 | 0.50 | 2 |
| Structure Distance (HD; average tie breaking) | 1436 | 0.90 | 0.50 | 0 |
| Structure Distance (HD; minimum tie breaking) | 1111 | 0.69 | 0.50 | 1 |
| Free Energy | 250 | 0.16 | 0.74 | 0 |
| Probability | 1554 | 0.97 | 0.52 | 4 |
| Ensemble Defect | 1527 | 0.95 | 0.51 | 2 |

**Table 2:** Results on synthetic structures of length 80. The “# Correct” is the number of correct solutions out of 1600. The “Correct Rate” is the ratio of the number of correct solutions and 1600. The “Unique Solver” column contains the number of structures for which a fitness function was the only fitness function to find a correct solution.

| Fitness Function | # Correct | Correct Rate | GC-percent | Unique Solver |
| --- | --- | --- | --- | --- |
| Structure Distance (BPD; arbitrary tie breaking) | 351 | 0.22 | 0.50 | 0 |
| Structure Distance (BPD; average tie breaking) | 454 | 0.28 | 0.50 | 1 |
| Structure Distance (BPD; minimum tie breaking) | 273 | 0.17 | 0.49 | 0 |
| Structure Distance (HD; arbitrary tie breaking) | 370 | 0.23 | 0.50 | 0 |
| Structure Distance (HD; average tie breaking) | 482 | 0.30 | 0.50 | 0 |
| Structure Distance (HD; minimum tie breaking) | 267 | 0.17 | 0.50 | 2 |
| Free Energy | 17 | 0.01 | 0.77 | 0 |
| Probability | 1201 | 0.75 | 0.59 | 112 |
| Ensemble Defect | 945 | 0.59 | 0.58 | 10 |

The second table contains results from the harder synthetic data set and shows probability with the highest correct rate and number of unique solutions. The same performance hierarchy is maintained, but probability pulls away by a larger margin. The number of unique solves is of particular interest where probability dominates all other fitness functions.

We tried several variations of structure distance. The results suggest that HD and BPD are comparable with perhaps a slight advantage to HD. Further, using an average to break ties is much better than arbitrary tie breaking (which is the norm in the literature) and minimum tie breaking.

### 3.2 Real RNAs

We consider the relative ranks of the true structure (or the closest analog) as a percentage. Consider an example percentage of 30.2%. This means, on average, the true structure was ranked at the position 30.2% below the highest rank. Higher ranks and therefore lower percentages are better as they correspond to the true structure being seen more favorably by the fitness function.

Probability achieved an mean relative rank of 21.2% with a standard deviation of 26.9% and a median of 8.0%. Ensemble defect achieved an mean relative rank of 48.0% with a standard deviation of 34.5% and a median of 47.8%. Structure distance achieved and mean relative rank of 48.2% with a standard deviation of 33.6% and a median of 47.9%.

These results are reflected in Figure 1 and Figure 2 where probability dominates the lower percentiles and ensemble defect and structure distance dominates the higher percentiles.

**Figure 1:**
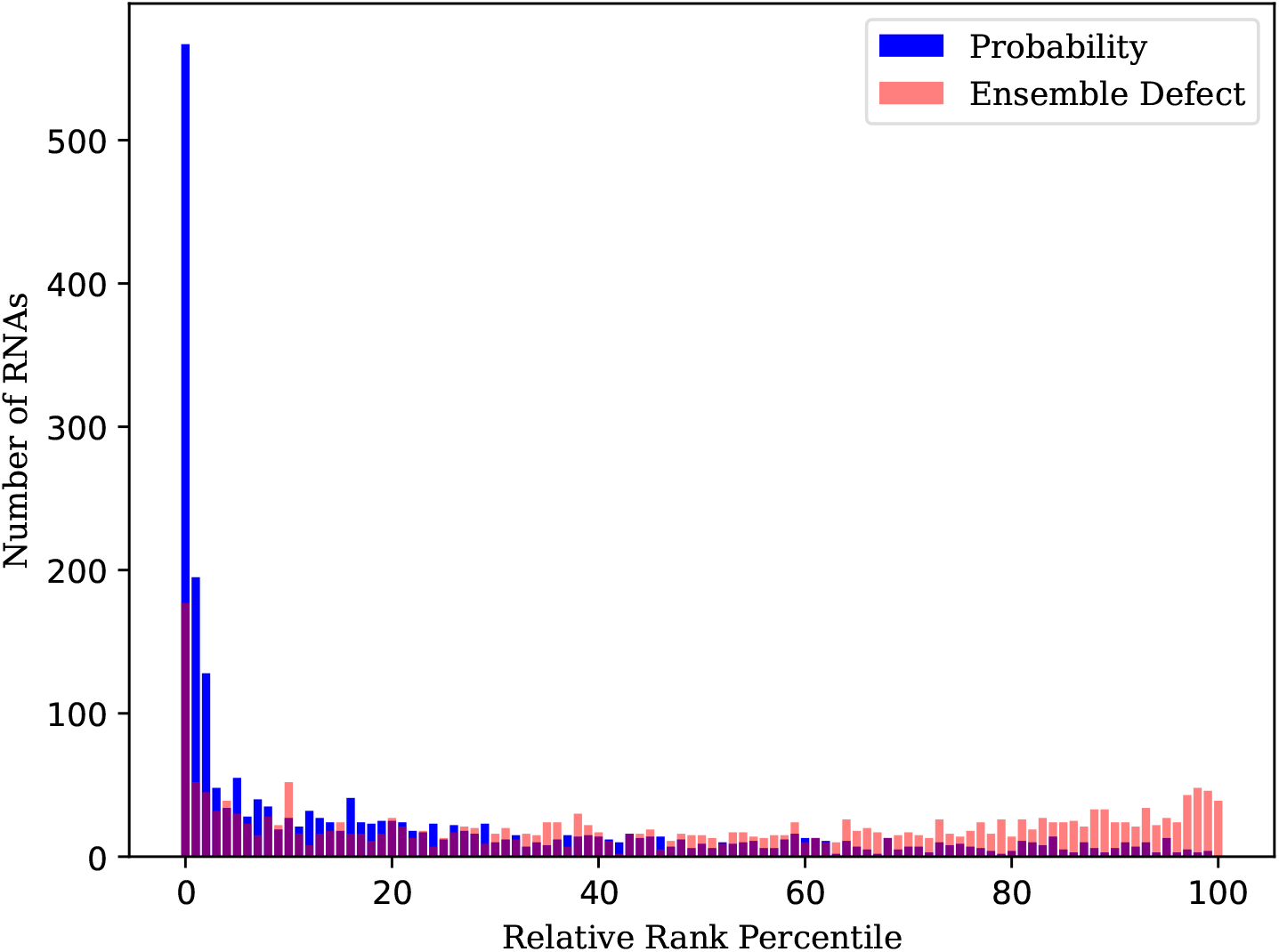
Comparison between probability and ensemble defect on real RNA sequences with known structures. Shows the number of RNAs for which the true structure (or the closest analog) fell into a certain percentile when ranked by a fitness function against other structures. Lower percentiles are better.

**Figure 2:**
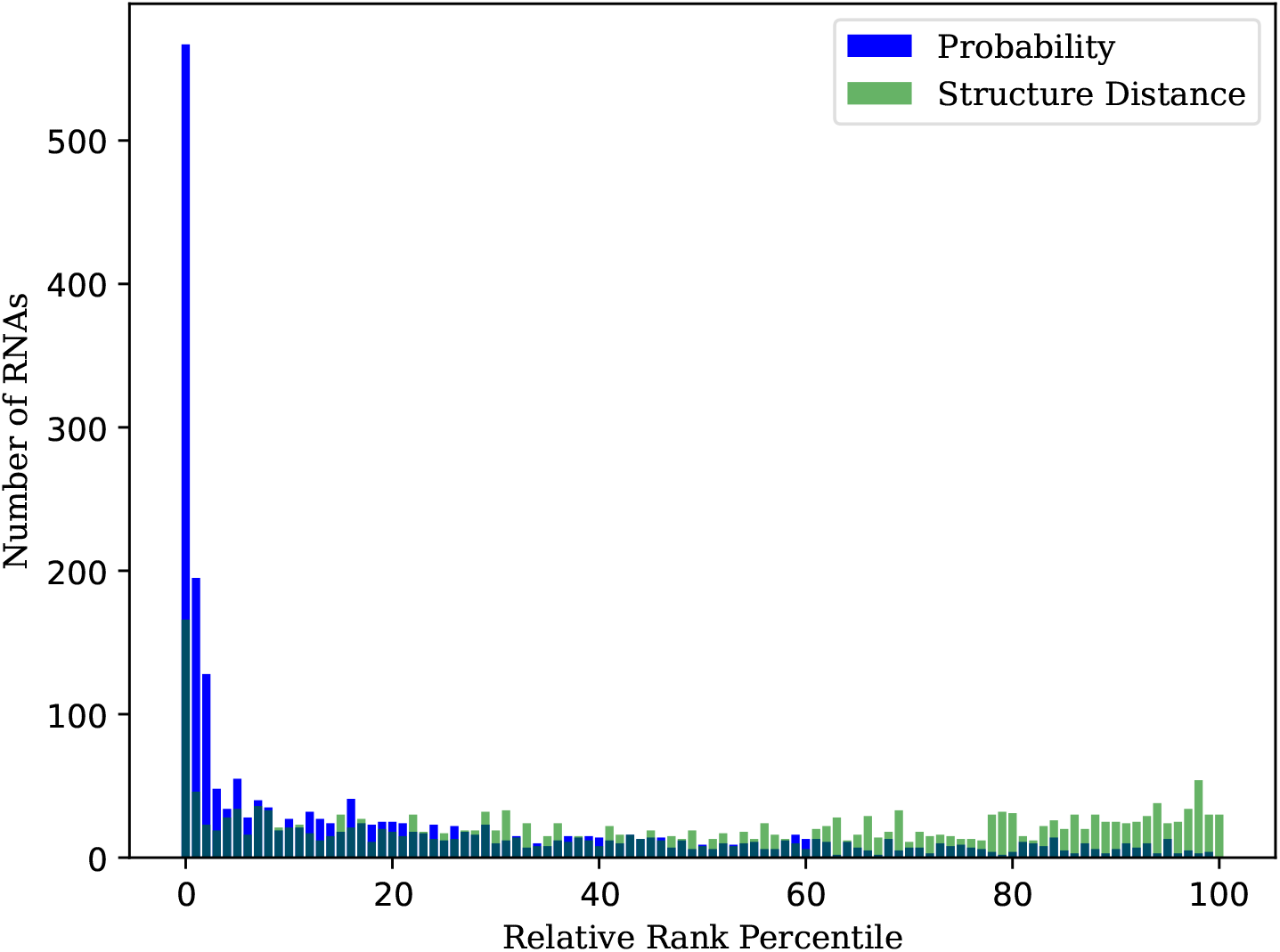
Comparison between probability and struture distance on real RNA sequences with known structures. Shows the number of RNAs for which the true structure (or the closest analog) fell into a certain percentile when ranked by a fitness function against other structures. Lower percentiles are better.

A representative example RNA is considered to illustrate why ensemble defect performs poorly compared to probability on natural RNAs. Consider the tRNA from Enterobacteria phage T5 with tRNAdb ID `tdbR00000022` [54]. Figure 3 shows a small but informative sample of structures from this RNA. The top two structures by probability rank are representative of all the top structures which either have a three-way multiloop (as in the first structure) or a four-way multiloop (as in the second structure). The true structure is well ranked by probability, because it is a variant of one these two structure clusters. The top ranked structure by ensemble defect is quite unnatural, containing an isolated pair. It is a compromise between these two different structural clusters and is indeterminate between three-way and four-way multiloops.

**Figure 3:**
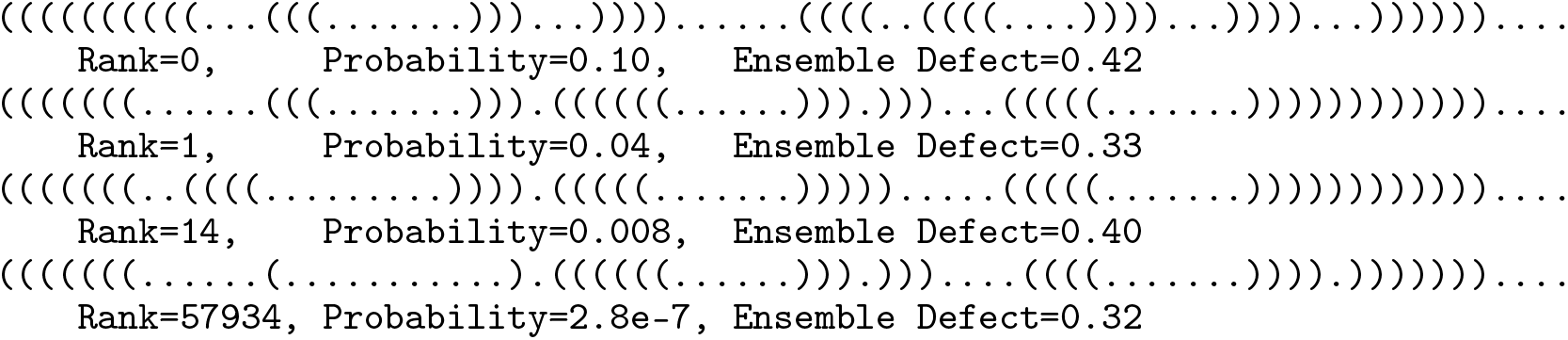
A sample of the structures for `tdbR00000022` in dot-bracket notation ranked by probability. In order, the top ranked structure, the second top ranked structure, the true structure, and top ranked structure by ensemble defect.

In addition, Figure 3 demonstrates a second separate failure mode that we observed for ensemble defect. Consider the second and third structures. Both are very similar, however the ensemble defect difference between them is large. This is because the first branch in the multiloop has been shifted in the 5^*′*^ direction, and a new pair has been added. The distance function employed by ensemble defect (see Equation (1)) is poor at capturing this kind of shifting. Note that the probability difference is similarly large in absolute terms, but probabilities do not have a flat distribution whereas ensemble defects have a flatter distribution, so the difference is less meaningful. This can be observed from the close ranks between the two structures when ordered by probability.

## 4 Discussion

Ensemble defect did not perform the best in our experiments. Probability performed better. Dirks *et al*. reported that probability and ensemble defect both performed well [47]. In their tests, both solved every RNA design challenge, with other fitness functions falling behind. We see a similar result in our smaller data set (Table 1), which we expect has a similar difficulty to that of Dirks *et al*. However, the larger data set (Table 2) shows probability having much stronger results. The explanation for the differences in our findings is that we tested on harder structures to design. If both fitness functions are solving every design challenge, then the challenges are not hard enough to properly distinguish the fitness functions.

We found the same pattern in real RNAs (Figure 1). Probability outperformed ensemble defect. An examination of specific RNAs where ensemble defect performs worse than probability is revealing. We see two major modes of failure for ensemble defect (Figure 3).

First, if there is more than one cluster of similar structures that are high probability, ensemble defect will compromise between the two clusters, which can lead to unrealistic averaging. We believe that a better interpretation when there are clusters is that a structure that is in at the center of a single cluster is likely, but one that is equidistant between two or more clusters is not.

Second, the distance function used in ensemble defect appears not to capture some important aspects of similarity. For example, a helix that moves in the 5’ or 3’ direction has a high defect compared to the original, since every base pair will be unaligned (like in Figure 3), however they are likely to be functionally similar structures.

It is possible that the issues with ensemble defect can be rectified by changing the distance function used (see Equation (1)). A more nuanced distance function addresses the second issue directly, and may have an indirect effect on the first. However, a primary strength of ensemble defect is that it can be efficiently computed. This is because of the intelligent choice of distance function [47]. It may be difficult to achieve such an efficient algorithm with a different distance function.

Table 1 and Table 2 record GC content. They suggest that free energy has a strong bias, which was expected given GC pairs have the lowest free energy change in the model. We expect to see 50% if the sequence was chosen randomly. Probability and ensemble defect appear to have a slight bias in GC content. The increase in GC content seems to be correlated with increased performance on the test as well as increased difficulty. This may be explained by the thermodynamic model predicting GC pairs are more stabilizing than other pairs. Thus, harder design challenges requiring must rely on using the most stabilizing pairs in some situations, increasing the required GC content. We note that Dirks *et al*. reported similar GC contents, albeit slightly elevated, possibly due to their data set having a bias [47].

Structure distance is the most widely used fitness function in the literature as we noted previously Section 1.2.1. The issue of minimum free energy tie breaking is not addressed in the existing literature. We address it here, and our results suggest that taking the average among the ties has the best result. We tested both base pair distance and structural Hamming distance. We did not observe any significant differences in performance, although structural Hamming distance may have a small advantage. Overall, no variant of structure distance came close to the level of probability. As such, we recommend that probability is preferred in the future over structure distance despite its popularity.

In Figure 2 we see that structure distance performs similarly to ensemble defect when used to rank real RNAs. However, it is less effective than ensemble defect on synthetic RNAs as seen in Table 2. This may be because the real RNA case exacerbates the disadvantages of ensemble defect. Future may investigate this further.

In the work presented here, we examined fitness functions in isolation. However, some existing techniques combine multiple fitness functions [31, 38]. Future work might test the performance of composite fitness functions.

Probability is the best performing fitness function, but it has a theoretical weakness described in Section 1.2.3; in short probabilities taken from different distributions may not be comparable. Future work may show that, in practice, RNA sequences of the same length have similar probability distributions over the corresponding structure ensemble. If this is true, then it would explain why comparing probabilities between different sequences works well in practice. Additionally, a better fitness function may be found that outperforms probability and does not have this issue.

In Section 1, we give three major components comprising an RNA design algorithm: the *computational model*, the *fitness function*, and the *search algorithm*. It is interesting that the choice of computational model informs the space of possible fitness functions. For example, probability and ensemble defect require that the model assigns the ensemble of structures probabilities. However, structure distance only requires a folding function. The choice of fitness function also constrains the search algorithm. To illustrate, [36] uses the properties of ensemble defect to weight mutations. A computational model based on the nearest neighbour thermodynamic model and dynamic programming recursions from [16] is widely assumed in RNA design, but there are alternatives [17]. We wonder if using a different paradigm altogether may affect the choice of fitness function and search algorithm.

## References

[1] Mark G Caprara and Timothy W Nilsen. RNA: versatility in form and function. Nature structural biology, 7(10):831–833, 2000.

[2] Harold S Bernhardt. The RNA world hypothesis: the worst theory of the early evolution of life (except for all the others). Biology direct, 7(1):1–10, 2012.

[3] Francis Crick. Central dogma of molecular biology. Nature, 227(5258):561–563, 1970.

[4] Jennifer A Doudna and Thomas R Cech. The chemical repertoire of natural ribozymes. Nature, 418(6894):222– 228, 2002.

[5] Alexander Serganov and Dinshaw J Patel. Ribozymes, riboswitches and beyond: regulation of gene expression without proteins. Nature Reviews Genetics, 8(10):776–790, 2007.

[6] Scott B Cohen, Mark E Graham, George O Lovrecz, Nicolai Bache, Phillip J Robinson, and Roger R Reddel. Protein composition of catalytically active human telomerase from immortal cells. Science, 315(5820):1850–1853, 2007.

[7] Ignacio Tinoco Jr and Carlos Bustamante. How RNA folds. Journal of molecular biology, 293(2):271–281, 1999.

[8] Colin B Reese. Oligo-and poly-nucleotides: 50 years of chemical synthesis. Organic & biomolecular chemistry, 3(21):3851–3868, 2005.

[9] Norbert Pardi, Michael J Hogan, Frederick W Porter, and Drew Weissman. mRNA vaccines—a new era in vaccinology. Nature reviews Drug discovery, 17(4):261–279, 2018.

[10] Joanna B Opalinska and Alan M Gewirtz. Nucleic-acid therapeutics: basic principles and recent applications. Nature Reviews Drug Discovery, 1(7):503–514, 2002.

[11] Farren J Isaacs, Daniel J Dwyer, Chunming Ding, Dmitri D Pervouchine, Charles R Cantor, and James J Collins. Engineered riboregulators enable post-transcriptional control of gene expression. Nature biotechnology, 22(7):841–847, 2004.

[12] James Chappell, Kyle E Watters, Melissa K Takahashi, and Julius B Lucks. A renaissance in RNA synthetic biology: new mechanisms, applications and tools for the future. Current opinion in chemical biology, 28:47–56, 2015.

[13] Ronny Lorenz, Stephan H Bernhart, Christian Höner zu Siederdissen, Hakim Tafer, Christoph Flamm, Peter F Stadler, and Ivo L Hofacker. ViennaRNA package 2.0. Algorithms for molecular biology, 6(1):1–14, 2011.

[14] Jessica S Reuter and David H Mathews. RNAstructure: software for RNA secondary structure prediction and analysis. BMC bioinformatics, 11(1):1–9, 2010.

[15] Liang Huang, He Zhang, Dezhong Deng, Kai Zhao, Kaibo Liu, David A Hendrix, and David H Mathews. Linear-fold: linear-time approximate RNA folding by 5’-to-3’dynamic programming and beam search. Bioinformatics, 35(14):i295–i304, 2019.

[16] Michael Zuker and Patrick Stiegler. Optimal computer folding of large RNA sequences using thermodynamics and auxiliary information. Nucleic acids research, 9(1):133–148, 1981.

[17] Elena Rivas. The four ingredients of single-sequence RNA secondary structure prediction. a unifying perspective. RNA biology, 10(7):1185–1196, 2013.

[18] Alexander Churkin, Matan Drory Retwitzer, Vladimir Reinharz, Yann Ponty, Jérôme Waldispühl, and Danny Barash. Design of RNAs: comparing programs for inverse RNA folding. Briefings in bioinformatics, 19(2):350– 358, 2018.

[19] Ivo L Hofacker, Walter Fontana, Peter F Stadler, L Sebastian Bonhoeffer, Manfred Tacker, and Peter Schuster. Fast folding and comparison of RNA secondary structures. Monatshefte für Chemie/Chemical Monthly, 125(2):167– 188, 1994.

[20] Manja Wachsmuth, Sven Findeiß, Nadine Weissheimer, Peter F Stadler, and Mario Mörl. De novo design of a synthetic riboswitch that regulates transcription termination. Nucleic acids research, 41(4):2541–2551, 2013.

[21] James Chappell, Melissa K Takahashi, and Julius B Lucks. Creating small transcription activating RNAs. Nature chemical biology, 11(3):214–220, 2015.

[22] Ivan Dotu, Juan Antonio Garcia-Martin, Betty L Slinger, Vinodh Mechery, Michelle M Meyer, and Peter Clote. Complete RNA inverse folding: computational design of functional hammerhead ribozymes. Nucleic acids research, 42(18):11752–11762, 2014.

[23] Hannah K Wayment-Steele, Do Soon Kim, Christian A Choe, John J Nicol, Roger Wellington-Oguri, Andrew M Watkins, R Andres Parra Sperberg, Po-Ssu Huang, Eterna Participants, and Rhiju Das. Theoretical basis for stabilizing messenger RNA through secondary structure design. Nucleic acids research, 49(18):10604–10617, 2021.

[24] Jeff Anderson-Lee, Eli Fisker, Vineet Kosaraju, Michelle Wu, Justin Kong, Jeehyung Lee, Minjae Lee, Mathew Zada, Adrien Treuille, and Rhiju Das. Principles for predicting RNA secondary structure design difficulty. Journal of molecular biology, 428(5):748–757, 2016.

[25] Michael Schnall-Levin, Leonid Chindelevitch, and Bonnie Berger. Inverting the Viterbi algorithm: an abstract framework for structure design. In Proceedings of the 25th international conference on Machine learning, pages 904–911, 2008.

[26] Édouard Bonnet, Pawel RzaŽewski, and Florian Sikora. Designing RNA secondary structures is hard. Journal of Computational Biology, 27(3):302–316, 2020.

[27] Jozef Haleš, Alice Héliou Ján Manúch, Yann Ponty, and Ladislav Stacho. Combinatorial RNA design: designability and structure-approximating algorithm in Watson–Crick and Nussinov–Jacobson energy models. Algorithmica, 79(3):835–856, 2017.

[28] Mirela Andronescu, Anthony P Fejes, Frank Hutter, Holger H Hoos, and Anne Condon. A new algorithm for RNA secondary structure design. Journal of molecular biology, 336(3):607–624, 2004.

[29] Anke Busch and Rolf Backofen. INFO-RNA–a fast approach to inverse RNA folding. Bioinformatics, 22(15):1823– 1831, 2006.

[30] Akito Taneda. MODENA: a multi-objective RNA inverse folding. Advances and applications in bioinformatics and chemistry: AABC, 4:1, 2011.

[31] Rune B Lyngsø, James WJ Anderson, Elena Sizikova, Amarendra Badugu, Tomas Hyland, and Jotun Hein. Frnakenstein: multiple target inverse RNA folding. BMC bioinformatics, 13(1):1–12, 2012.

[32] Álvaro Rubio-Largo, Leonardo Vanneschi, Mauro Castelli, and Miguel A Vega-Rodríguez. Multiobjective metaheuristic to design RNA sequences. IEEE Transactions on Evolutionary Computation, 23(1):156–169, 2018.

[33] Juan Antonio Garcia-Martin, Peter Clote, and Ivan Dotu. RNAiFOLD: a constraint programming algorithm for RNA inverse folding and molecular design. Journal of bioinformatics and computational biology, 11(02):1350001, 2013.

[34] Gerard Minuesa, Cristina Alsina, Juan Antonio Garcia-Martin, Juan Carlos Oliveros, and Ivan Dotu. MoiRNAiFold: a novel tool for complex in silico RNA design. Nucleic acids research, 49(9):4934–4943, 2021.

[35] Sinem Sav, David JD Hampson, and Herbert H Tsang. SIMARD: A simulated annealing based RNA design algorithm with quality pre-selection strategies. In 2016 IEEE Symposium Series on Computational Intelligence (SSCI), pages 1–8. IEEE, 2016.

[36] Joseph N Zadeh, Brian R Wolfe, and Niles A Pierce. Nucleic acid sequence design via efficient ensemble defect optimization. Journal of computational chemistry, 32(3):439–452, 2011.

[37] Stanislav Bellaousov, Mohammad Kayedkhordeh, Raymond J Peterson, and David H Mathews. Accelerated RNA secondary structure design using preselected sequences for helices and loops. RNA, 24(11):1555–1567, 2018.

[38] Fernando Portela. An unexpectedly effective Monte Carlo technique for the RNA inverse folding problem. BioRxiv, page 345587, 2018.

[39] Xiufeng Yang, Kazuki Yoshizoe, Akito Taneda, and Koji Tsuda. RNA inverse folding using Monte Carlo tree search. BMC bioinformatics, 18(1):1–12, 2017.

[40] Tristan Cazenave and Thomas Fournier. Monte Carlo inverse folding. In Monte Carlo Search International Workshop, pages 84–99. Springer, 2020.

[41] Robert Kleinkauf, Martin Mann, and Rolf Backofen. antaRNA: ant colony-based RNA sequence design. Bioinformatics, 31(19):3114–3121, 2015.

[42] Jeehyung Lee, Wipapat Kladwang, Minjae Lee, Daniel Cantu, Martin Azizyan, Hanjoo Kim, Alex Limpaecher, Snehal Gaikwad, Sungroh Yoon, Adrien Treuille, et al. RNA design rules from a massive open laboratory. Proceedings of the National Academy of Sciences, 111(6):2122–2127, 2014.

[43] Rohan V Koodli, Benjamin Keep, Katherine R Coppess, Fernando Portela, Eterna participants, and Rhiju Das. EternaBrain: Automated RNA design through move sets and strategies from an internet-scale rna videogame. PLoS computational biology, 15(6):e1007059, 2019.

[44] Peter Eastman, Jade Shi, Bharath Ramsundar, and Vijay S Pande. Solving the RNA design problem with reinforcement learning. PLoS computational biology, 14(6):e1006176, 2018.

[45] Frederic Runge, Danny Stoll, Stefan Falkner, and Frank Hutter. Learning to design RNA. arXiv preprint 1812.11951, 2018.

[46] Douglas H Turner and David H Mathews. NNDB: the nearest neighbor parameter database for predicting stability of nucleic acid secondary structure. Nucleic acids research, 38(Suppl_1):D280–D282, 2010.

[47] Robert M Dirks, Milo Lin, Erik Winfree, and Niles A Pierce. Paradigms for computational nucleic acid design. Nucleic acids research, 32(4):1392–1403, 2004.

[48] Rune B Lyngsø, Michael Zuker, and Christian NS Pedersen. Internal loops in RNA secondary structure prediction. In Proceedings of the third annual international conference on Computational molecular biology, pages 260–267, 1999.

[49] Hamid Dadkhahi, Jesus Rios, Karthikeyan Shanmugam, and Payel Das. Fourier representations for black-box optimization over categorical variables, 2022.

[50] John S McCaskill. The equilibrium partition function and base pair binding probabilities for RNA secondary structure. Biopolymers: Original Research on Biomolecules, 29(6-7):1105–1119, 1990.

[51] Joseph N Zadeh, Conrad D Steenberg, Justin S Bois, Brian R Wolfe, Marshall B Pierce, Asif R Khan, Robert M Dirks, and Niles A Pierce. NUPACK: Analysis and design of nucleic acid systems. Journal of computational chemistry, 32(1):170–173, 2011.

[52] Stefan Wuchty, Walter Fontana, Ivo L Hofacker, and Peter Schuster. Complete suboptimal folding of RNA and the stability of secondary structures. Biopolymers: Original Research on Biomolecules, 49(2):145–165, 1999.

[53] Max Ward, Amitava Datta, Michael Wise, and David H Mathews. Advanced multi-loop algorithms for RNA secondary structure prediction reveal that the simplest model is best. Nucleic acids research, 45(14):8541–8550, 2017.

[54] Frank Jühling, Mario Mörl, Roland K Hartmann, Mathias Sprinzl, Peter F Stadler, and Joern Pütz. tRNAdb 2009: compilation of tRNA sequences and tRNA genes. Nucleic acids research, 37(Suppl_1):D159–D162, 2009.

